# METATRYP v 2.0: Metaproteomic Least Common Ancestor Analysis for Taxonomic Inference Using Specialized Sequence Assemblies - Standalone Software and Web Servers for Marine Microorganisms and Coronaviruses

**DOI:** 10.1101/2020.05.20.107490

**Authors:** Jaclyn K. Saunders, David Gaylord, Noelle Held, Nick Symmonds, Chris Dupont, Adam Shepherd, Danie Kinkade, Mak A. Saito

## Abstract

We present METATRYP version-2 software that identifies shared peptides across organisms within environmental metaproteomics studies to enable accurate taxonomic attribution of peptides during protein inference. Improvements include: ingestion of complex sequence assembly data categories (metagenomic and metatranscriptomic assemblies, single cell amplified genomes, and metagenome assembled genomes), prediction of the Least Common Ancestor (LCA) for a peptide shared across multiple organisms, increased performance through updates to the backend architecture, and development of a web portal (https://metatryp.whoi.edu). Major expansion of the marine database confirms low occurrence of shared tryptic peptides among disparate marine microorganisms, implying tractability for targeted metaproteomics. METATRYP was designed for ocean metaproteomics and has been integrated into the Ocean Protein Portal (https://oceanproteinportal.org); however, it can be readily applied to other domains. We describe the rapid deployment of a coronavirus-specific web portal (https://metatryp-coronavirus.whoi.edu/) to aid in use of proteomics on coronavirus research during the ongoing pandemic. A Coronavirus-focused METATRYP database identified potential SARS-CoV-2 peptide biomarkers and indicated very few shared tryptic peptides between SARS-CoV-2 and other disparate taxa, sharing 0.1% peptides or less (1 peptide) with the Influenza A & B pan-proteomes, establishing that taxonomic specificity is achievable using tryptic peptide-based proteomic diagnostic approaches.

**Statement of significance:** When assigning taxonomic attribution in bottom-up metaproteomics, the potential for shared tryptic peptides among organisms in mixed communities should be considered. The software program METATRYP v 2 and associated interactive web portals enables users to identify the frequency of shared tryptic peptides among taxonomic groups and evaluate the occurrence of specific tryptic peptides within complex communities. METATRYP facilitates phyloproteomic studies of taxonomic groups and supports the identification and evaluation of potential metaproteomic biomarkers.

## Introduction

In metaproteomics the mixture of a large number of organisms within each sample collected from a natural environment creates challenges in the attribution of peptides to specific proteins. This is especially problematic in instances where exact tryptic peptide sequences are shared between two or more organisms. This potential for shared peptides across proteins can create uncertainty in protein inference and taxonomic attribution. In bottom-up proteomics, the primary method used in metaproteomics to date, whole proteins are typically digested into smaller peptides with the enzyme trypsin. Since bottom-up metaproteomics directly measures these short tryptic peptides, as opposed to entire protein sequences, it is essential to understand the degree of shared peptides across proteins and taxonomic groups when assigning attributes of diverse environmental communities. Previously, we described the development of the METATRYP software which evaluates multiple organisms for shared peptides [1]. METATRYP takes the full predicted proteome of an organism based on its reference genome, performs an *in silico* tryptic digestion of the proteins, and then stores the tryptic peptides of that organism within a single SQL database. Multiple taxa proteomes are stored within the SQL database. Using METATRYP tools, the database can be queried to identify how many taxa share a specific peptide (or list of peptides), and it can also identify the total number of specific tryptic peptides shared across multiple organisms and for other phyloproteomic analyses. The former application has aided in the development of targeted metaproteomic biomarkers for assessing environmental changes in space or time [1-4]. A useful result of the latter application was the observation that the percentage of shared peptides between distinct marine microbial taxa was low, often in the single digit percentages, implying that the design of biomarker targets for species or even subspecies level analyses was tractable if sufficient care was taken.

In this manuscript, we describe version 2 of the METATRYP software (https://github.com/WHOIGit/metatryp-2.0). We have added additional features to improve its usability and performance. A major improvement was the addition of new data categories for different sequencing assembly methods, specifically those associated with assembled metagenomic and metatranscriptomic data as well as single cell amplified genomes (SAGs). METATRYP v 2 now supports these three specific data categories: “Genomes” for reference cultured isolates, “Specialized assemblies” from SAGs and MAGs, and “Meta-omic assemblies” from metagenomic and metatranscriptomic assemblies. This greatly expands the utility of METATRYP since cultured genomes are often unavailable from natural environmental populations due to many organisms being difficult to culture with classical microbiological techniques [5], or being only recently identified taxa. As a result, the availability of single cell genomes amplified and sequenced from the ocean environment (SAGs), metagenome assembled genomes (MAGs), and assembled metagenomics and metatranscriptomic data can contribute greatly to the identification and interpretation of metaproteomic data. Yet because these metagenomic and metatranscriptomic resources have varying levels of completeness and confidence in their functional and taxonomic assignments, maintaining them as separate categories of tryptic peptides within the database structure is particularly useful. In addition, METATRYP v 2 now supports the calculation of a Least Common Ancestor (LCA) among shared tryptic peptides. For comparison, the Unipept web portal [6] has some similar functionality in identifying shared tryptic peptides and interpreting the Least Common Ancestor; however, Unipept relies on the Uniprot database which does not incorporate the wealth of environmental meta-omic sequencing available. Also, the Unipept portal does not support local curated database construction where users can evaluate unpublished sequencing resources like those of newly sequenced organismal genomes or novel environmental sequencing not yet available in the Uniprot curated database whereas METATRYP can be installed locally for use with custom curated databases. The addition of these new sequence assembly data categories enables better prediction of shared peptides through enhanced representation of environmental sequence variability. The METATRYP marine web portal currently contains a total of 182,354,079 unique peptides from 19,104,353 submitted protein sequences combined across all three data categories.

Multiple improvements to the METATRYP software architecture and additional features were added to v 2. In order to improve performance speed and support these larger data categories (especially, metagenomic and metatranscriptomic assemblies), the METATRYP SQL backend was converted from SQLite in METATRYP v 1 to a PostgreSQL backend in METATRYP v 2. Additional software and PostgreSQL implementations support the LCA analysis. METATRYP v 2 uses the same tryptic digest rules applied to METATRYP v 1, following trypsin-based digestion rules for proteins with peptides 6-22 amino acids in length [1]. Here we describe the technical improvements within METATRYP v 2, then demonstrate how metagenomics resources allow increased understanding of Least Common Ancestor interpretations of metaproteomic results. Additionally, we provide an overview of the METATRYP web portal for marine microorganisms and the rapid deployment of a coronavirus-specific METATRYP web portal demonstrating the application of METATRYP to various research fields. Finally, a private API was added providing LCA analysis functionality to the Ocean Protein Portal [7], enabling METATRYP to be inserted into other pipelines in the future.

## Implementation

### Database Backend Upgrades

METATRYP is built upon a Relational Database Management System (RDMS). Version 1 was built using a SQLite database. While this database management system was sufficient for single reference genomes, it was lacking in speed needed for expanded sequencing data categories. In order to increase the speed of database construction (ingestion of proteomes), database searching, and expanded functionality, we have upgraded the database management system to a PostgreSQL backend which is an object-relational database. The Postgres backend has provided improvements in speed, as well as enhanced flexibility in searching.

### Support for New Data Categories

The advent of environmental *de novo* sequencing and assembly has identified an entire realm of microorganisms previously unknown to the scientific world, as many environmental microorganisms are not readily isolated using classical microbiological techniques [5]. METATRYP v 1 focused on the construction of a search database using reference organismal genomes, METATYRP v 2 is capable of handling newer types of sequencing and assemblies thus opening the search space to a much greater range of organisms likely found within the environment of interest. Within the field of metaproteomics, there has been great emphasis placed on the need for curated and appropriate search databases to be utilized for peptide-to-spectrum matching (PSM) [8, 9]; this also holds true for the construction of a METATRYP database for evaluation of shared tryptic peptides in an environmental sample. METATRYP relies upon protein sequences predicted from genomic sequencing and does not currently take into account any post-translational modifications (PTMs).

In order to expand the environmentally-relevant search space, METATRYP v 2 now handles newer assemblies from sources like metagenome and metatranscriptome assemblies, metagenome-assembled-genomes (MAGs), and single cell amplified genomes (SAGs). The incorporation of these newer data categories, in addition to the traditional single organism reference genome, greatly expands the environmental variability (and therefore, potential for shared tryptic peptides) within an environmental sample. In order to manage the larger sequencing databases generated by incorporation of this environmental data, and thus the greater burden of a larger search space, improvements were made to the back-end search database (see section “Database Back-end Upgrades”). The database schema (Figure 1) for METATRYP was expanded to not only include these different data categories, but to identify them as separate search spaces as the uncertainty of taxonomic identity among the sequencing categories is variable and should be taken into consideration during interpretation. The new data categories roughly mirror the original reference genome schema. However, additional tables are required to properly map meta-omic data as multiple taxa are contained within a single meta-omic assembly file.

**Figure 1.**
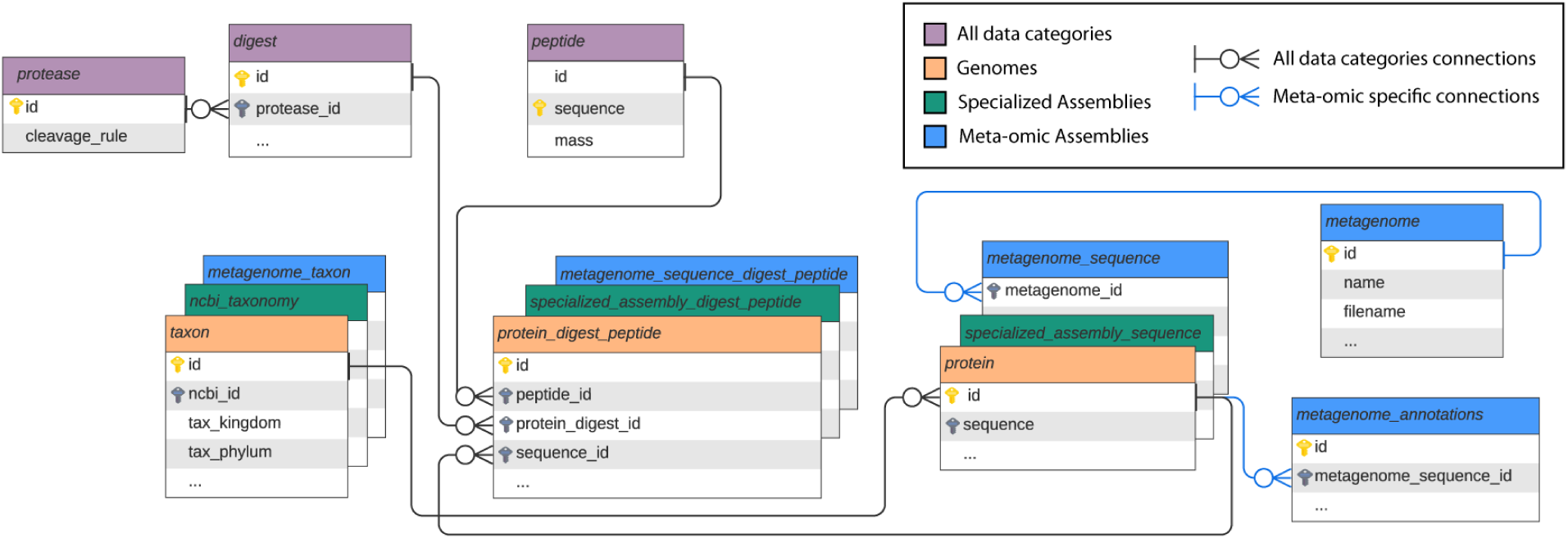
Core elements of the METATRYP v 2 database schema. An entity relationship diagram depicting the core tables for the three sequencing categories (orange tables: “Genome”, green tables: “Specialized Assembly”, and blue tables: “Meta-omic Assembly” data categories). The purple tables are shared tables among all three data categories containing information about the tryptic digestion rules (*protease* and *digest* tables) as well as the amalgamation of all unique tryptic peptide sequences found across all three data categories (*peptide*). The lines connecting the tables represent links between the data tables. The three data categories are stacked where tables represent similar information for each category. The metagenome data category requires two additional data tables as there are multiple taxa stored within a single meta-omic assembly sequencing file which requires an additional *metagenome_annotations* table for parsing; the blue connecting lines represent meta-omic specific data linkages.

Environmental sequencing data is also more likely to have more frequent occurrences of ambiguous bases in assemblies. These are base locations where it is uncertain what the correct amino acid should be, sometimes a result of low-quality base calling by the sequencing technology or due to a single location where there are multiple possibilities for the amino acid at that single location in the assembly that cannot be determined. METATRYP v 2 will recognize ambiguous bases, specifically the base symbol “X”, which represents the presence of an unknown or ambiguous amino acid during the ingestion phase. As the specific amino acid represented by “X” is unknown, the exact tryptic peptide cannot be predicted. During ingestion, METATRYP will identify proteins which contain an “X” and report to stdout the affected protein. If there is a tryptic peptide containing the “X”, that peptide will not be included in the METATRYP peptide table; however, all other tryptic peptides from that protein will be included in the peptide table.

### Least Common Ancestor Analysis

Sequence homology is conserved among more closely related organisms. However, it is possible that tryptic peptides, especially shorter ones around 6 amino acids, may occur by chance across multiple taxa without a direct shared ancestry. In order to identify the occurrence of shared tryptic peptides either through shared evolutionary history or through stochastic variance in sequence, we have added Least Common Ancestor (LCA) analysis to METATRYP v 2. The LCA analysis incorporates the phylogenetic lineage of the sequences imported into the METATRYP databases (Figure 1), then calculates the “Least Common Ancestor” by finding the unifying phylogenetic point for all the organisms containing the tryptic peptide queried. In order for METATRYP to identify the common point in the taxonomic lineage, it requires a consistent taxonomic lineage to be used across the database for each proteome submitted. For METATRYP the shared phylogeny of the taxonomic groups is identified by pulling the taxonomic lineages from the National Center for Biotechnology Information (NCBI) Taxonomy Database [10]. For the creation of a user-generated METATRYP database capable of LCA analysis, the user can submit the NCBI taxon id number (taxid) for the input sequence files, and METATRYP will pull the taxonomic lineage information for each organism using Biopython [11] and Pandas [12] libraries in Python 3. This lineage information is then used to calculate the LCA among the organisms with shared peptides via the PostgreSQL Longest Common Ancestor function, which METATRYP uses to return the LCA for each sequencing data category.

### Web Portal & API

A primary goal of releasing the METATRYP software originally was to enable other users to create and curate customized databases for searching tryptic peptides, specifically with a focus on marine microbial communities. METATRYP v 2 expands on this goal through the creation of a web server and API which can be queried easily by users, without the need to install and run the software locally. The METATRYP v 2 site can be found at https://metatryp.whoi.edu/. This web server takes as input a peptide sequence, multiple peptides, or a full protein sequence submitted into a text box by a user which is then *in silico* digested into tryptic peptides. METATRYP then searches for the occurrence of these peptides across three different marine-specific data categories: an organismal reference genome data category (“genomes”) same as METATRYP v 1 (SI Table 1), and new data categories for “specialized assemblies” (SI Table 2) which currently contains 4,783 Archaeal and Bacterial MAGs [13, 14] assembled by binning metagenomic sequences [15-17] from the TARA Oceans sequencing project [18], and a metagenomic & metatranscriptomic assembly data category (“meta-omic assembly”) [3, 4, 19] (SI Table 3) which currently contains 4,863,985 predicted proteins. Ideally, additional MAGs and SAGs will be added to the METATRYP web portal database in an effort to broaden taxonomic coverage in the marine environment. The addition of Eukaryotic SAGs [20] would significantly extend the diversity of the current database.

Results from a METATRYP query are then returned to the user in an interactive drop-down table, showing the presence of the peptides within these data categories and the LCA result for each category (Figure 2). This example shows the results for three different peptides that have been searched simultaneously (LSHQAIAEAIGSTR, VNSVIDAIAEAAK, VAAEAVLSMTK) which are used for the default search on the web portal if a user does not enter a sequence query. To expand the search results for each peptide, the user clicks on the peptide sequence link; shown here are the results for peptide VNSVIDAIAEAAK in the “Genome” and “Meta-omic” data categories. Displayed are the taxa and their associated NCBI Taxonomy ID numbers (taxid) which link out to that taxon’s entry within the NCBI Taxonomy Database. For the “Genome” category, the Joint Genome Institute (JGI) Integrated Microbial Genomes & Microbiomes (IMG) genome IDs are also shown, where available, with links out to IMG as the predicted proteomes for all those in the “Genome” category were collected from in the current version of this database [21].

**Figure 2.**
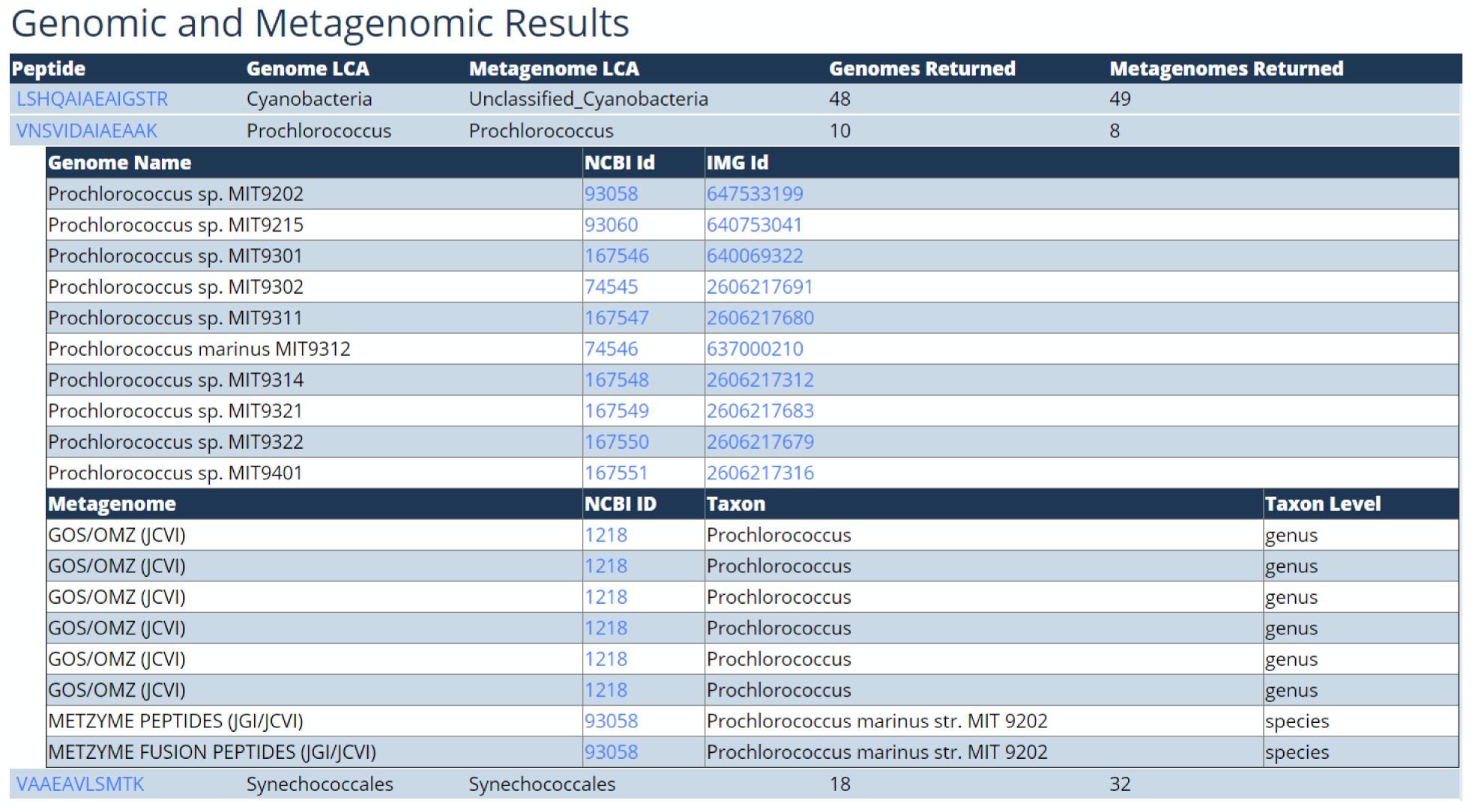
Genomic and Meta-omic query results from the marine METATRYP web portal for the peptides LSHQAIAEAIGSTR, VNSVIDAIAEAAK, VAAEAVLSMTK. The LCA results for these three separate peptides indicate the varying degrees of taxonomic uniqueness among the peptides in the “Genome” and “Meta-omic Assembly” data categories. In these two data categories, LSHQAIAEAIGSTR is unique to the Phylum Cyanobacteria. VNSVIDAIAEAAK is unique to the genus *Prochlorococcus*. VAAEAVLSMTK is unique to the Order Synechococcales. All of these peptides show potential as biomarkers at these varying taxonomic levels according to evaluation by the “Genome” and “Meta-omic Assembly” databases.

METATRYP can also compare entire organismal proteomes to identify the frequency of shared peptides across taxa within a given sequencing data category. Those more familiar with nucleic acid sequencing often incorrectly imagine that because translation to amino acid space results in loss of variable DNA codon information, peptides will lack the ability to taxonomically resolve species or subspecies. This feature previously existed in METATRYP v 1 for generating peptide redundancy tables within the “Genome” sequencing data category and was used to show the relatively low occurrence of shared peptides across disparate taxa in the open ocean microbiome [1]. This feature can now be implemented within METATRYP v 2 on all three major data categories: “Genome”, “Meta-omic Assemblies”, and “Specialized Assemblies”. Within the web portal, there is now a visualization tool for creating ordered heatmaps of shared peptide frequencies among taxa for the genome and meta-omic data categories. This visualization page, “Peptide Redundancy Heatmaps”, was built in Python 3.7 using the Jupyter environment [22], Pandas [12], and Seaborn [23]. Users can select what taxa they wish to compare within a given data category, and a heatmap is generated and displayed on the page below in a .png format. Displayed in the heatmaps are the percentage of pairwise shared peptides between taxa in a specified data category where the percentage is calculated as the number of shared peptides between taxon A and taxon B divided by the total number of peptides in taxon A. Given the varying levels of genome completeness for a specific taxon in the “Specialized Assembly” data category, this percentage should be viewed with more caution. Due to the aggregate nature of meta-omic assemblies, where many taxa of highly variable coverage depth are present within each dataset, this heatmap visualization feature is not currently supported in the web portal for this data category.

### Example of Rapid Deployment: Coronavirus Domain Application

The capabilities of the METATRYP software make it applicable to scientific domains outside of the marine microbial ecology and biogeochemistry fields. The ability to identify shared peptides in metaproteomics is critical to other metaproteomics studies of mixed communities, especially in the development of biomarkers for targeted metaproteomics. An example application of METATRYP v 2 to other domains is the creation of a Coronavirus-focused database and the rapid deployment of a Coronavirus-focused METARTYP web portal (https://metatryp-coronavirus.whoi.edu). The database for this web portal uses the predicted proteomes for multiple of Riboviruses with a focus on Betacoronaviruses, including SARS-Cov-2 responsible for the COVID-19 outbreak [24], SARS-CoV-1 strains like SARS strain Frankfurt 1 [25] isolated during the 2003-2004 SARS outbreak, Middle East Respiratory syndrome-related (MERS) Coronavirus strains [26], and strains associated with the common cold [27] like Human Coronavirus strains NL63 [28], HKU1 [29] and 229E [30] for a total of 94 Coronavirus taxa in the database. It also contains the human proteome, the African Green Monkey (*Chlorocebus aethiops sabaeus*) proteome as it is the taxonomic source of the Vero cell line commonly used in virus replication studies and plaque assays [31], common oral bacteria [32], six *Lactobacillus* strains associated with the human microbiome [33], the most common Influenza strains (Influenza A: H1N1 & H3N2; Influenza B) as well as other Influenza strains, and common proteomic contaminants in the CRAPome [34]. All taxa included in the Coronavirus database and their associated NCBI Taxonomy IDs (SI Table 4) are listed on the databases page in the web portal (https://metatryp-coronavirus.whoi.edu/database). In order to capture the variability of sequences, we pulled all the proteins (aside from those in the CRAPome) for each taxon from the NCBI Identical Protein Groups (IPG) Database using taxon sequence identifiers (SI Table 4). The IPG Database enables collection of a single non-redundant entry for each protein translation found from several sources at NCBI, including annotated coding regions in GenBank and RefSeq, as well as records from SwissProt and PDB [35]. One sequence for each identical protein group was collected for NCBI taxa with >= 10 identical protein groups, collecting proteins from the specified taxon and all its children from the NCBI Taxonomy Database. However, for the taxon “Severe acute respiratory syndrome-related coronavirus” (NCBI txid: 694009), only protein groups from that specific txid level were recruited (using the flag “txid694009[Organism:noexp]” in the query). This taxon should only contain SARS-CoV-1 related proteins; however, it may contain some non-SARS-CoV-1 sequences due to inconsistent nomenclature during the emergence of this relatively new pathogen. Proteins were collected using Biopython [11] and the NCBI Entrez [36] E-Utilities API [37].

By using the Identical Protein Groups, the database captures sequence variability while reducing redundancy in database construction. This non-redundant collection of protein sequences per taxon is essentially the collection of the known pan-proteomes for each taxon; it is the representation of the predicted proteome of a single organism’s genome plus all the sequence variation captured from sampling a population of organisms within a taxonomic group. For example, the taxon *Homo sapiens* pan-proteome from IPG has >1,000,000 proteins capturing sequence variability from the population of human sequences in the NCBI database, whereas an individual human genome contains <20,0000 protein coding genes [38]. Due to the varying sequencing efforts of among some taxa, the length of the proteomes in this database may vary. For example, the taxon Severe acute respiratory syndrome coronavirus 2 (SARS-CoV-2, 2019-nCov, COVID-19 virus; taxid 2697049) has 1,752 unique Identical Protein Groups whereas the taxon Bat coronavirus isolate RaTG13 (taxid 2709072), SARS-CoV-2’s closest sequenced animal virus precursor [39], has 10 unique Identical Protein Groups as of this writing (May 3, 2020) even though the genomes of these viruses are roughly the same length. Since the population of SARS-CoV-2 has been sequenced more frequently, more sequence variability has been captured in the NCBI databases. Due to the collection of the pan-proteomes for the Coronavirus METATRYP database, the peptide redundancy heatmaps need to be viewed with caution, as taxa which have received a higher degree of sampling effort will have more peptides associated with them. Therefore, it is important to take into consideration both combinations of pairwise taxa comparisons for heatmap calculations (both sides of the heatmap separated by the diagonal). For example, one should evaluate the percentage of shared peptides between SARS-CoV-2 and RaTG13 where the total number of tryptic peptides for SARS-CoV-2 is in the denominator and also where the total number of tryptic peptides in the denominator is for RaTG13. When calculating peptide redundancy between organisms, METATRYP reports this calculation as “individual percent”.

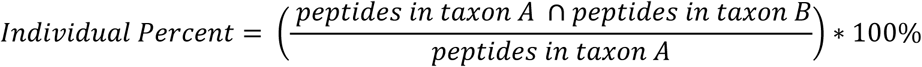

The percentage of shared peptides across taxa can also be calculated by the number of peptides shared between taxa with the combined total number of peptides for both taxa in the denominator, METATRYP reports this as “union percent”.

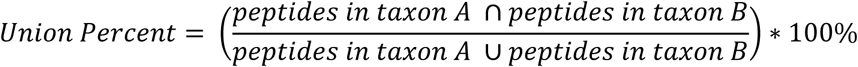

However, this also needs to be interpreted with care as when comparing a taxon that may have only been sequenced once (few total peptides) with a broadly sampled taxon (many peptides due to environmental variability), the signal of the rarely sampled taxon may be reduced due to the large n of the heavily sequenced organism. In addition, viewing “union percents” when comparing an organisms with a small proteome vs an organism with a large proteome would also skew any signal of shared peptides, for example SARS-CoV-2 only encodes 12 proteins/genome [24], whereas the human genome encodes ∼120,000 proteins. Even though the NCBI IPG Database is not used in the marine METATRYP web portal, this same effect may be observed when comparing organisms with highly uneven proteome sizes, say if marine phages are added to the database, or with taxa that have varying levels of genome sequencing completeness, such as with Specialized Assemblies like MAGs and SAGS.

## Results & Discussion

### Backend Upgrades for METATRYP v 2 Provide Improved Performance

The switch from a SQLite database backend in METATRYP v 1 to a PostgreSQL database backend has resulted in significant improvements in performance and functionality of METATRYP. In particular, this transition has resulted in improved computational times and facilitated the addition of LCA analyses to the software package. To test clock times, a database was constructed based upon the reference genomes from 136 marine microbial taxa (SI Table 5) in both versions of METATRYP. SI Table 6 shows the benchmarks associated with the construction of these genome-only databases for comparison, with METATRYP v 2 taking only 17% of the time it took METATRYP v 1 to construct the same database. By converting to a PostgreSQL backend, the computational time to query the example database with the example peptide “LSHQAIAEAIGSTR” was reduced over 10,000x, dropping from 1442.87 seconds using METATRYP v 1 to 0.11 seconds with METATRYP v 2. The CPU utilization for METATRYP v 2 was also lower than for v 1.

It is noted that setting up a PostgreSQL server on a local machine requires administrative permissions and is more complex than SQLite which is more user-friendly and requires fewer system dependencies. Therefore, for users without local administrative permissions, METATRYP v 1 with its SQLite backend remains a good lighter weight option (albeit with slower performance and without the capacity to handle diverse data categories and LCA analyses). The PostgreSQL backend facilitates the LCA analysis through a call to the PostgreSQL Longest Common Ancestor function. In order to facilitate ease of setup for the marine microbial research community, we have added a copy of this pre-constructed marine microbial METATRYP v 1 SQLite database to the METATRYP v 1 code repository (https://github.com/saitomics/metatryp). A complete pre-constructed marine microbial database for METATRYP v 2 containing sequence data for 136 genomes (SI Table 5) and the base schema for all three data categories is included in the METATRYP v 2 code repository (https://github.com/WHOIGit/metatryp-2.0).

### Use of New Data Categories in METATRYP for Marine Microbial Communities

Using the results of the peptides set as the default example search in the METATRYP web portal (LSHQAIAEAIGSTR, VNSVIDAIAEAAK, VAAEAVLSMTK), we see the results for these three peptides have varying levels of taxonomic LCAs, ranging from Species-to Phylum-levels of taxonomic specificity (Figures 2 & 3). For peptide VNSVIDAIAEAAK, the LCA across all data categories is the genus indicating *Prochlorococcus*, indicating that this peptide appears unique to this genus in the marine microbial community and is therefore a potential biomarker for targeted metaproteomics that allows species level specificity. For peptide VAAEAVLSMTK, the LCA for all categories is the Order Synechococcales as this peptide is found within the Genus *Prochlorococcus* and its sister Genus *Synechococcus*.

Taxonomic assignment of the original sequences of predicted proteins for the different data categories ranges in uncertainty. From reference genomes from cultured isolates being the most certain, to metatranscriptomic & metagenomic assemblies providing the least certainty in taxonomic assignment of the source proteins. The “Specialized assemblies” of SAGs and MAGs exist somewhere in between on this taxonomic assignment uncertainty spectrum. Peptide LSHQAIAEAIGSTR (Figure 3) demonstrates this level of uncertainty within the “Meta-omic assembly” category as a source protein for this peptide in the GOS/OMZ (JCVI) metagenome cannot be identified with below the Phylum level of Cyanobacteria (Figure 2) and in the “Specialized assembly” category containing thousands of MAGs, this peptide is found in 94 Cyanobacterial MAGs and one Verrumicrobial MAG (Verrucomicrobiales_bacterium_strain_NP1000), resulting in a LCA of “Bacteria”. Notably, this peptide is from the Global Nitrogen Transcriptional Regulation Protein (NtcA) which is highly conserved across the Cyanobacteria [40]. While this peptide has been previously identified as a potential biomarker for Cyanobacteria [1], the addition of the “Specialized Assembly” data category identified the possible presence of this peptide in another phyla warranting further investigation. Interestingly, upon further investigation, the protein in MAG Verrucomicrobiales_bacterium_strain_NP1000, identified from metagenomes in the Red Sea [14], is >99% identical to an NtcA in the Cyanobacterium genus *Synechococcus* (NCBI accession WP_067098506.1). This Verrumicrobial *ntcA* may occur within this genome as a result of horizontal gene transfer or as an artifact of the MAG binning process. Either way, it indicates that peptide LSHQAIAEAIGSTR should be used with caution as a Cyanobacterial biomarker in environments with abundant Verrumicrobia. However, this is an unlikely scenario as Cyanobacteria tend to be far more abundant in marine environments than Verrumicrobia. Query results from a full sequence NtcA from *Prochlorococcus* MED4 (NCBI GenBank accession CAE18705.1) show that other peptides, such as “LVSFLMVLCR”, may be more appropriate if targeting *Prochlorococcus* only (SI Figure 1). By separating sequence types into different categories, METATRYP allows the user to balance the varying levels in confidence of taxonomic attribution, where reference genomes from cultured isolates are the best in taxonomic quality but more incomplete in environmental coverage, and vice versa for environmental sequences.

**Figure 3.**
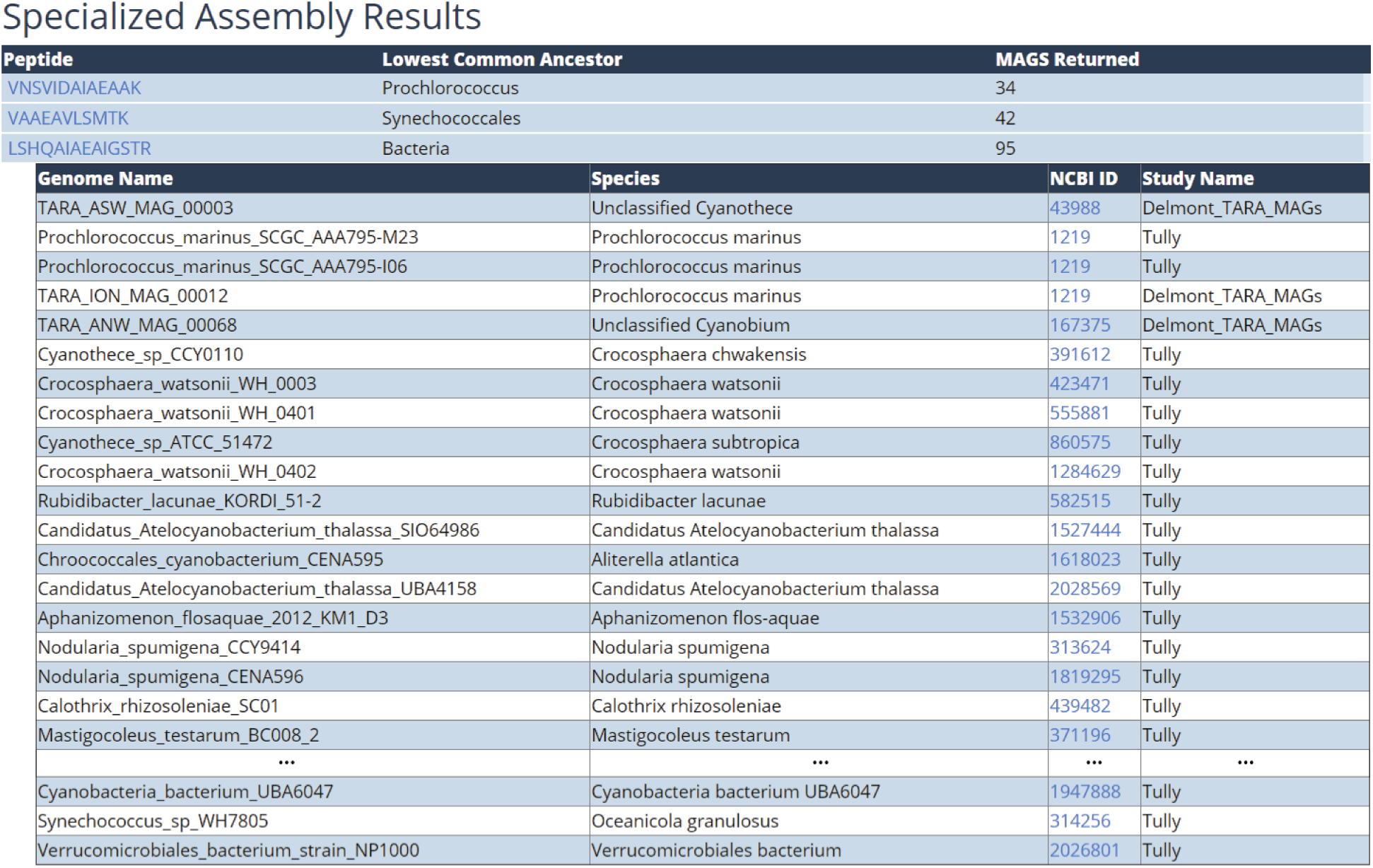
Specialized Assembly abbreviated query results from the marine METATRYP web portal for the peptides LSHQAIAEAIGSTR, VNSVIDAIAEAAK, VAAEAVLSMTK. Among the MAGs currently in the METATRYP web portal database, VNSVIDAIAEAAK & VAAEAVLSMTK are unique to MAGs at the Genus level for *Prochlorococcus* and *Synechococcus*, respectively. However, the peptide LSHQAIAEAIGSTR, while indicating specificity at the Phylum level of Cyanobacteria in the “Genome” and “Meta-omic Assembly” data categories, reports 94 Cyanobacterial MAGs with this peptide and 1 Verrumicrobia MAG with this peptide, warranting further investigation of this peptide as a potential Cyanobacterial biomarker and considerations for use of this potential biomarker in environments which may have abundant Verrumicrobial populations.

The addition of 4,783 MAGs to METATRYP has provided further insight into the prevalence of shared peptides across taxonomic groups in marine microbial communities. An analysis with METATRYP v 1 using a database of 51 single reference genomes from common pelagic marine microorganisms demonstrated very little overlap in shared peptides across different taxonomic groups [1]. Expanding this analysis to the 4,783 MAGs shows a similar pattern of a very low occurrence of shared tryptic peptides across disparate taxa. Figure 4 shows the individual percentages of shared peptides across a random selection of 25 MAGs. In general, taxa share <1% of tryptic peptides, with a few clusters of more closely related organisms sharing more tryptic peptides -- for example, Gammaproteobacterial and Euryarchaeotal clusters highlighted with red outlines -- where the taxa share between 1-2% or 2-14% of tryptic peptides in each cluster, respectively. A similar pattern is shown when the cross-wise comparison of taxa is expanded to 100 MAGs (SI Figures 2 & 3). These results demonstrate that there should be sufficient resolution to discern between taxa using tryptic peptides identified by metaproteomic analyses, especially when coupled to LCA analysis tools like METATRYP to confirm peptide taxonomic origin.

**Figure 4.**
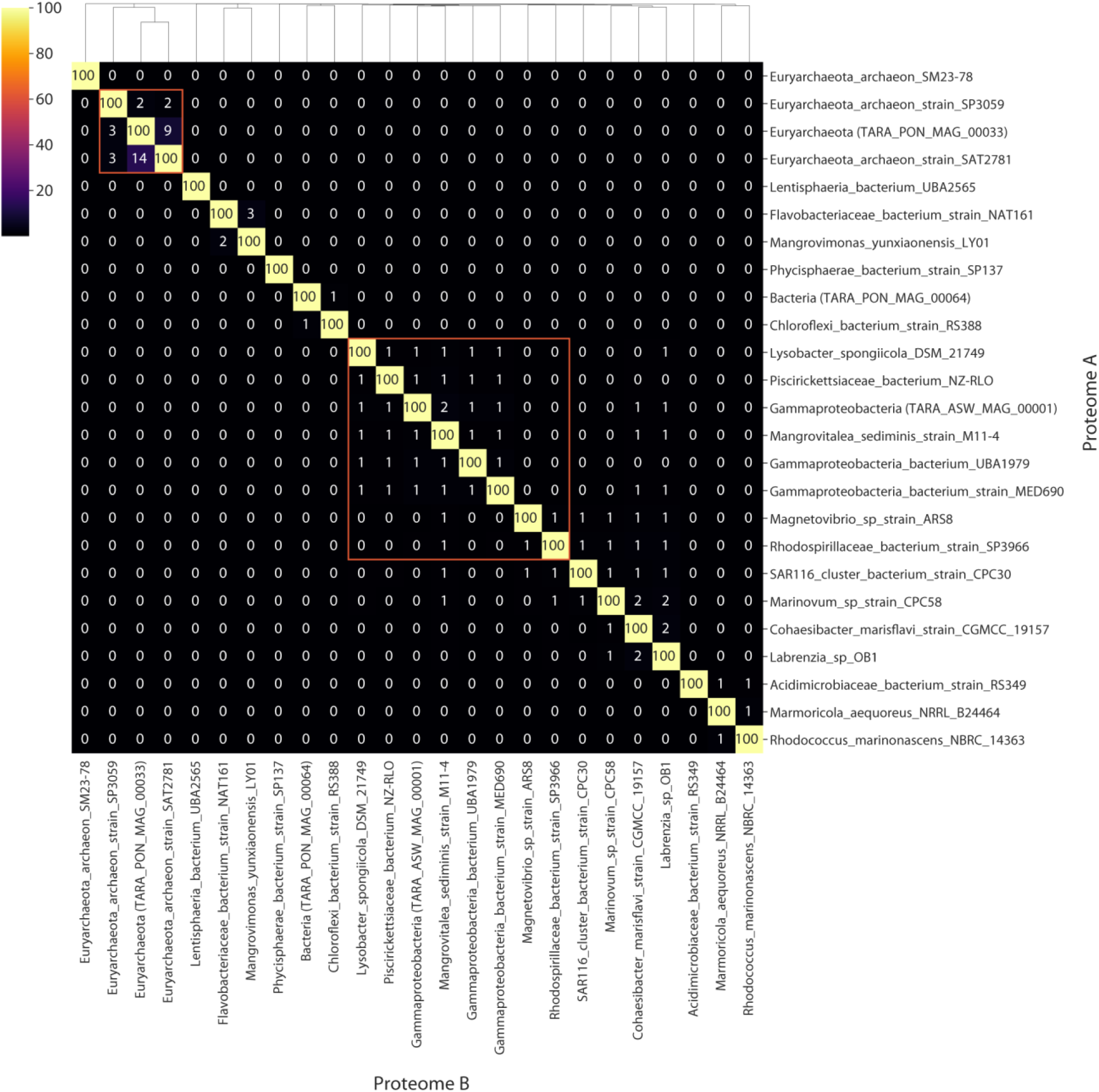
Heatmap with cluster dendrogram displaying the individual percents of shared tryptic peptides across 25 randomly selected MAGs from the new “Specialized Assembly” data category. The individual percent is calculated as the percentage of shared peptides across Proteome A and Proteome B divided by the number of total peptides in Proteome A. The data here show a very low frequency of shared tryptic peptides across this random sample of MAGs (SI Figures 2 & 3 show 100 randomly selected MAGs displaying a similar trend). The red outlines highlight two example regions of Euryarchaeotal and Gammaproteobacterial MAGs showing some overlap in shared tryptic peptides among these taxonomic groups.

### Example of Rapid Deployment: Coronavirus Domain Application

Analysis of the Coronavirus-focused METATRYP instance is similar to the observations of marine-focused METATRYP where there is a rather low frequency of shared peptides across disparate taxa with a higher frequency of shared peptides across more closely related taxa (Figure 5; SI Figures 4 & 5). Within the broader group of the 81 taxa associated with Severe acute respiratory syndrome (SI Table 4; SI Figures 6 & 7), there is a greater occurrence of shared tryptic peptides. Notably, there is a large cluster of strains associated with the 2003-2004 SARS-CoV-1 outbreak (such as strain Human Coronavirus Frankfurt 1) that cluster together with >90% shared tryptic peptides among the strains. SARS-CoV-2 responsible for the ongoing COVID-19 pandemic is found in a separate cluster among these taxa, clustering most closely with bat-hosted Coronaviruses with 84% shared peptides from the SARS-CoV-2 group with the bat RaTG13 virus. The strain SARS Coronavirus Frankfurt 1 (SARS-CoV-1) shares up to 35% of tryptic peptides with the SARS-CoV-2 group. SARS-CoV-2 shares 1% or less of shared tryptic peptides with other taxonomic groups outside the Severe acute respiratory syndrome related Coronaviruses (Figure 5; SI Figures 6 & 7). This analysis of shared tryptic peptides highlights that SARS-CoV-2 is indeed different, in tryptic peptide space, from other viral pathogens like Influenza. Only 1 peptide is shared between SARS-CoV-2 and the major Influenza subtypes, sharing peptide “DGQAYVR” from chain C of a SARS-CoV-2 spike glycoprotein [41]. The reference genome for SARS-CoV-2 (NCBI Reference Sequence NC_045512.2) contains 12 protein coding genes with 828 tryptic peptides, thus a SARS-CoV-2 genome shares 0.1% or less tryptic peptides with the Influenza pan-proteomes. Thus, differentiating SARS-CoV-2 from other major viral pathogens is tractable using proteomic analyses. Users developing diagnostic assays should take care to independently confirm their informatic analyses, as the METATRYP Coronavirus instance is intended as a software example and the current database may not be maintained concurrently with the emergence of new sequencing data.

**Figure 5.**
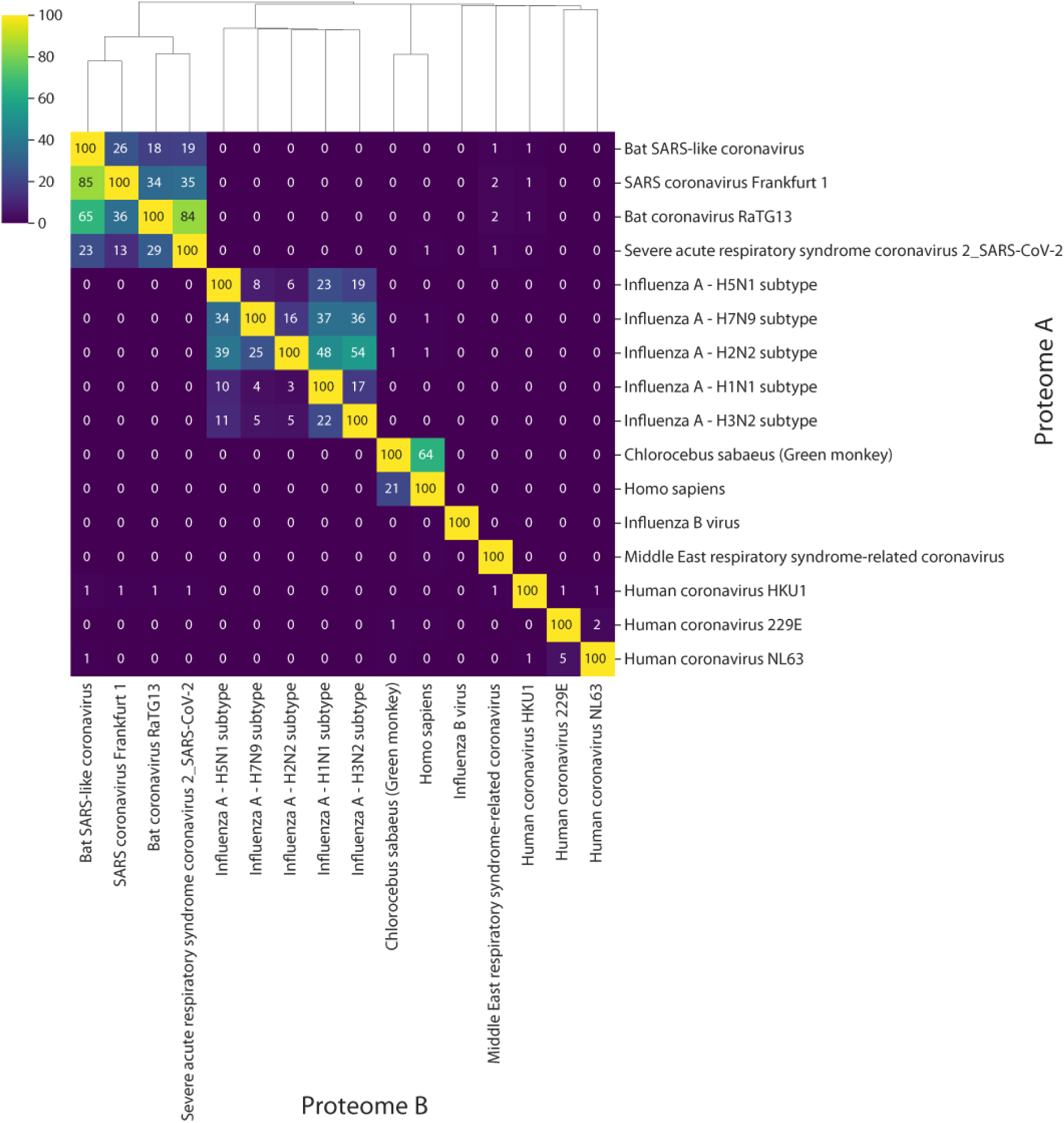
Heatmap with cluster dendrogram displaying shared tryptic peptides among a subsample of the taxa in the METATRYP Coronavirus web portal database. In general, there is a very low occurrence of shared tryptic peptides (<1%) across disparate taxa. Higher frequencies of shared tryptic peptides are shown among more closely related taxonomic groups. Severe acute respiratory syndrome-related Coronaviruses form a cluster in the top left corner. MERS-related coronavirus shares <1% of tryptic peptides with all taxa depicted here. The “common cold” strains of Coronavirus (HKU1, 229E, and NL63) form a separate cluster in the bottom right corner. These Coronavirus clusters are distinct and very different from the Influenza A cluster in the middles, as the Severe acute respiratory syndrome-related viruses share <1% of shared tryptic peptides with Influenza A and B. *Homo sapiens* and *Chlorcebus sabaeus* form a distinct group, sharing more tryptic peptides with each other than with any other taxonomic groups.

The Coronavirus-focused METATRYP database enables the investigation of shared tryptic peptides across multiple key taxa. A METATRYP database applied in this domain may help identify biomarker peptides that could be used in quantitative proteomics to identify SARS-Cov-2-specific peptides in metaproteomic samples, whether those samples come from human test subjects with complex oral microbiomes or from environmental samples. Using a spike glycoprotein sequence encoded by the S gene from SARS-CoV-2 (NCBI accession YP_009724390.1) as a query, multiple potential SARS-CoV-2 peptides are identified as potential biomarkers for the virus. This protein contains 69 tryptic peptides, with 17 of those peptides being specific to the taxon SARS-CoV-2. Some of the peptides are found across multiple taxa: 52 peptides appear in SARS-CoV-2 and at least one other taxon. Among these, there are peptides which appear to be more conserved across the Severe acute respiratory syndrome related viruses: 16 peptides are found in 65 or more Severe acute respiratory related viral taxa (ranging in viruses that infect humans to other hosts like bats, civets, and pigs). The peptide ‘GIYQTSNFR’ may be a potential biomarker for this group of viruses in general, as it is found in all 81 taxa from the Severe acute respiratory syndrome related virus group, but not found in other taxa within the database. Given its applications in metaproteomics, METATRYP may be a powerful tool for proteomics applied to SARS-CoV-2 wastewater-based epidemiology used to track community spread of COVID-19 infections [42, 43], as a combination of the Coronavirus-specific database with other environmental databases (like marine microbial METATRYP) may provide insight into potential tryptic peptide biomarkers in sewage effluent.

## Conclusions

This manuscript announces the release of version 2 of the METATRYP software package for assessing shared tryptic peptides in complex communities. In addition to the standalone software, we have created web portals for METATRYP v 2 instances that use specialized databases for the marine microbial research community as well as the coronavirus research community. A private API for the marine microbial instance of METATRYP supports LCA analysis in the Ocean Protein Portal [7] and may be further developed into a public API which can be connect to automated pipelines in the future, like GalaxyP [44]. This major release of METATRYP features an upgraded SQL backend that supports faster speeds for database construction and data queries, enables maintenance of separate sequencing data categories within the database, and facilitates LCA analysis of shared peptides across taxa. The main METATRYP web portal database consists of three data categories: reference genomes, specialized sequencing assemblies, and meta-omic sequencing assemblies. Users can readily query these databases for the occurrence of specific peptide sequences and visualize the frequency of shared peptides across taxa in the reference genomes and specialized assemblies. The expansion of METATRYP beyond reference genomes allows for more complete coverage of the diversity found in environmental communities; however, these newer sequencing assembly types also carry higher degrees of uncertainty in the taxonomic attributions assigned to the source proteins. The major scientific findings of Saito et al. (2015), that redundancy of tryptic peptides across disparate taxa is rare, is supported when these new broader sequencing data categories were included in the search database suggesting taxonomic specificity of the majority of tryptic peptides. METATRYP aids in the selection of biomarker peptides for identification of specific taxonomic groups at varying taxonomic levels. We also demonstrated how METATRYP can be applied to proteomics analyses in other scientific domains through the creation of the METATRYP Coronavirus web portal. Users can query the occurrence of shared peptides encoded by various coronavirus genomes and other relevant taxa. Using this portal, we showed that the SARS-CoV-2 Coronavirus has the most shared tryptic peptides with its closest bat precursor virus, has some shared peptides with SARS-CoV-1, and is very different from the “common flu”. METATRYP is a flexible software package to assess taxonomic occurrence of shared peptides applicable to proteomics studies of complex systems valuable for the identification of biomarkers and phyloproteomic analysis of complex communities.

## Acknowledgements

We would like to thank A. Murat Eren, Tom Delmont, Ben Tully, Elaina Graham, and John Heidelberg for graciously providing MAG sequences and additional taxonomic information facilitating incorporation into the METATRYP database. This work was made possible by grants from the National Science Foundation EarthCube Data Infrastructure Grant NSF-ICER 1639714 and Division of Ocean Science grants NSF-OCE 1657766 & 1924554 and the Gordon and Betty Moore Foundation grants 8453 and 3782. JKS was additionally supported by a NASA Postdoctoral Program Fellowship. METATRYP v 2 is a product of the Ocean Protein Portal (OPP). The OPP team is a collaboration between the Saito laboratory, the Information Services Application group, and the Biological and Chemical Oceanography Data Management Office all at the Woods Hole Oceanographic Institution.

