## Supplemental Figures for "METATRYP v 2.0: Metaproteomic Least Common Ancestor Analysis for Taxonomic Inference Using Specialized Sequence Assemblies - Standalone Software and Web Servers for Marine Microorganisms and Coronaviruses"

### Specialized Assembly Results

| Peptide | Lowest Common Ancestor | MAGS Returned |
|---|---|---|
| MSPASR | cellular_organisms | 39 |
| LTPQPSGQNTINTIGER | Prochlorococcus | 1 |
| TLMEVIK | Prochlorococcus | 33 |
| GLEGASTEMVER | Synechococcales | 10 |
| TIFFPGDPAER | Bacteria | 88 |
| VYESGEEITVALLR | Bacteria | 61 |
| ENSLFGVLSLLTGHR | Bacteria | 53 |
| FYHAIAFTR | Synechococcales | 16 |
| VEMITAPANSVLR | Prochlorococcus | 33 |
| AIEADASVG | None | 0 |
| LLLLQGLSSR | None | 0 |
| ILQTETMIETLTHR | Bacteria | 58 |
| LVSFLMVLCR | Prochlorococcus | 38 |
| DFGVASEK | Prochlorococcus | 11 |
| GITIDLR | Bacteria | 41 |
| LSHQAIAEAIGSTR | Bacteria | 95 |
| LLGDLK | cellular_organisms | 115 |
| DSGLLTIER | Prochlorococcus | 4 |
| ITVFDPIALSK | Prochlorococcus | 2 |

**SI Figure 1.** Specialized Assembly query results from a full sequence NtcA from *Prochlorococcus* MED4 (NCBI GenBank accession CAE18705.1). Results show that other peptides, such as "LVSFLMVLCR", may be more appropriate if targeting the Genus *Prochlorococcus*, specifically.

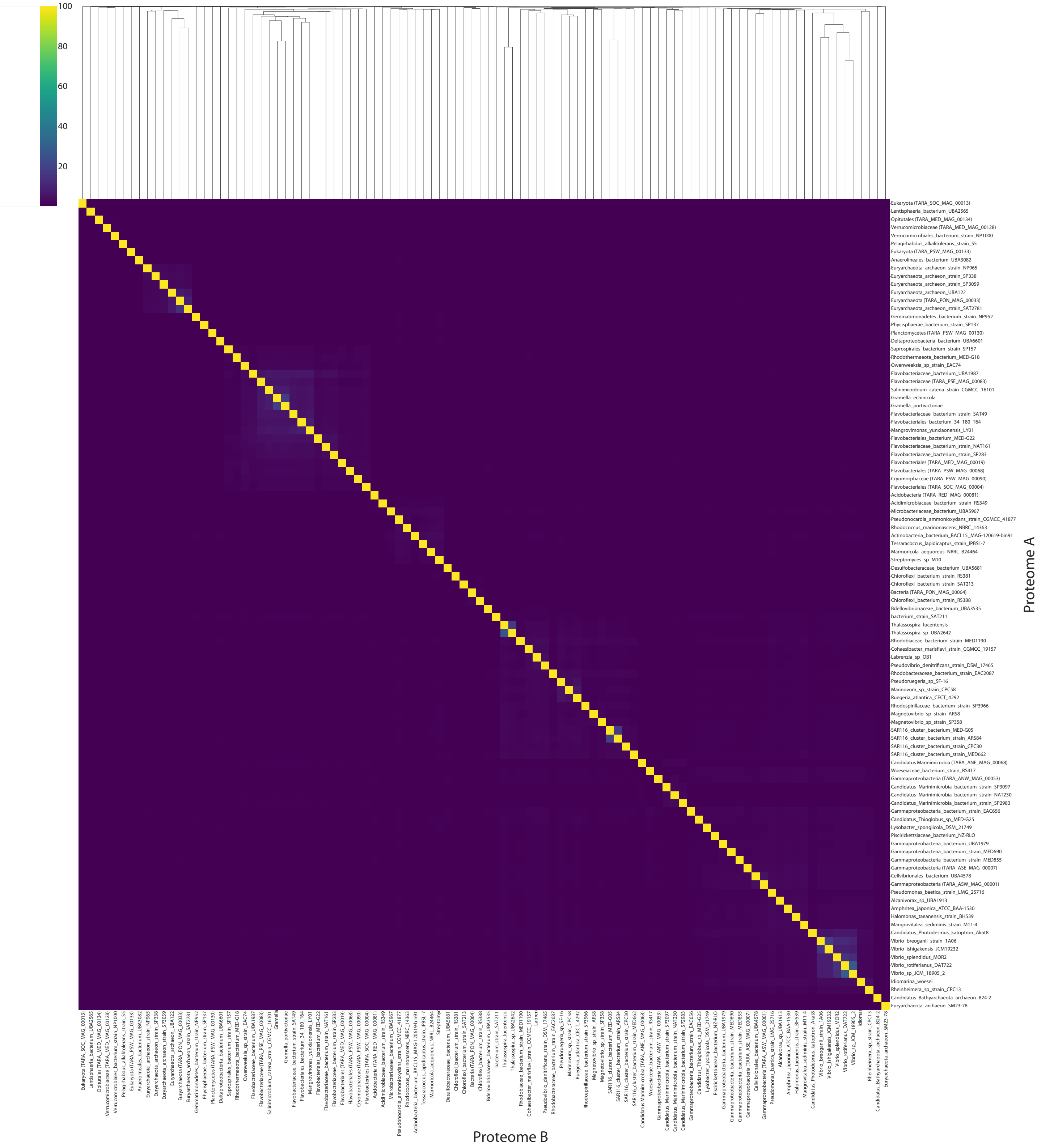


**SI Figure 2.** Heatmap with cluster dendrogram displaying the individual percents of shared tryptic peptides across 100 randomly selected MAGs from the new "Specialized Assembly" data category (see SI Figure 3 for annotations).

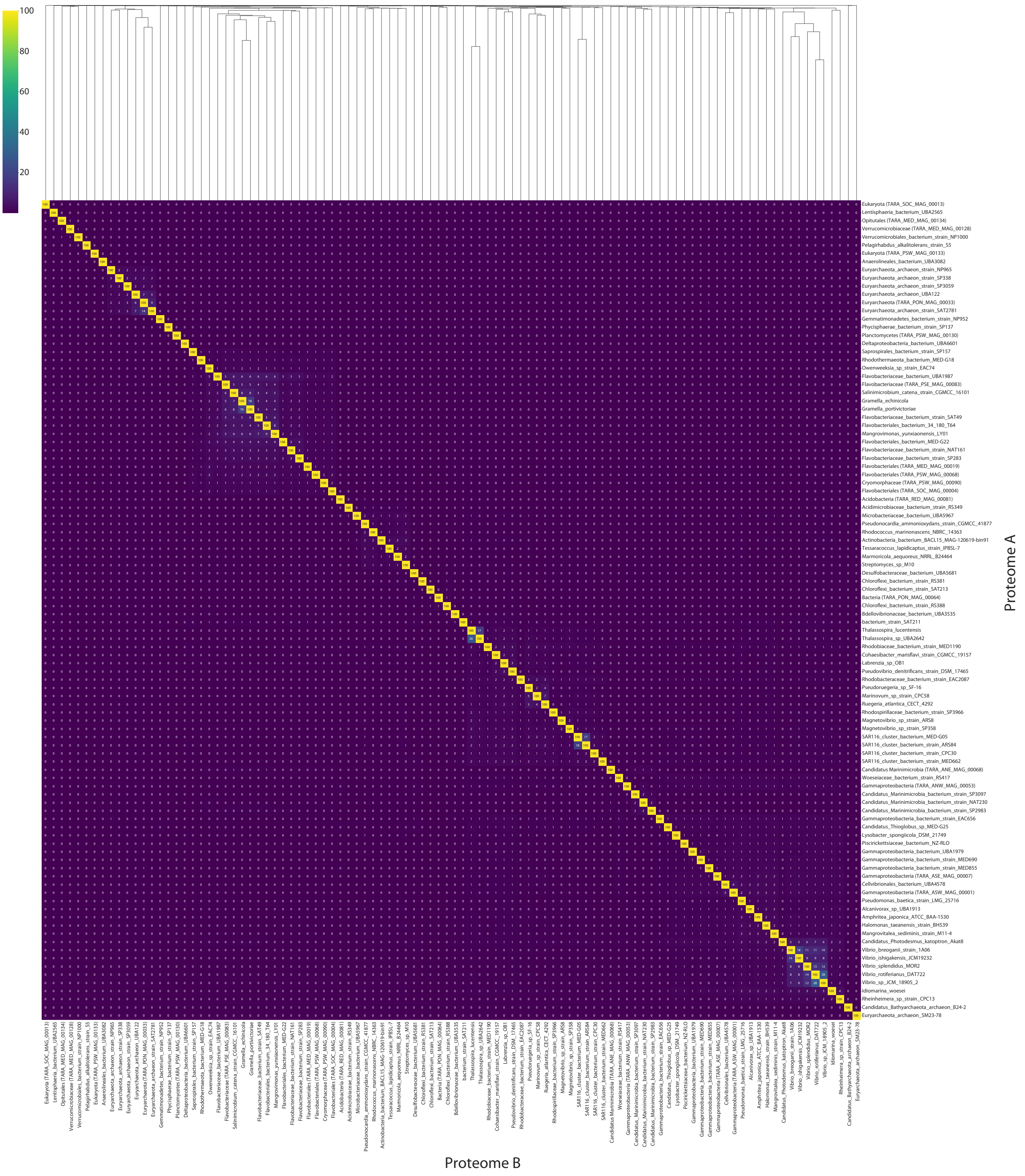


**SI Figure 3.** Heatmap with cluster dendrogram displaying the individual percents of shared tryptic peptides across 100 randomly selected MAGs from the new "Specialized Assembly" data category with annotated percentages.

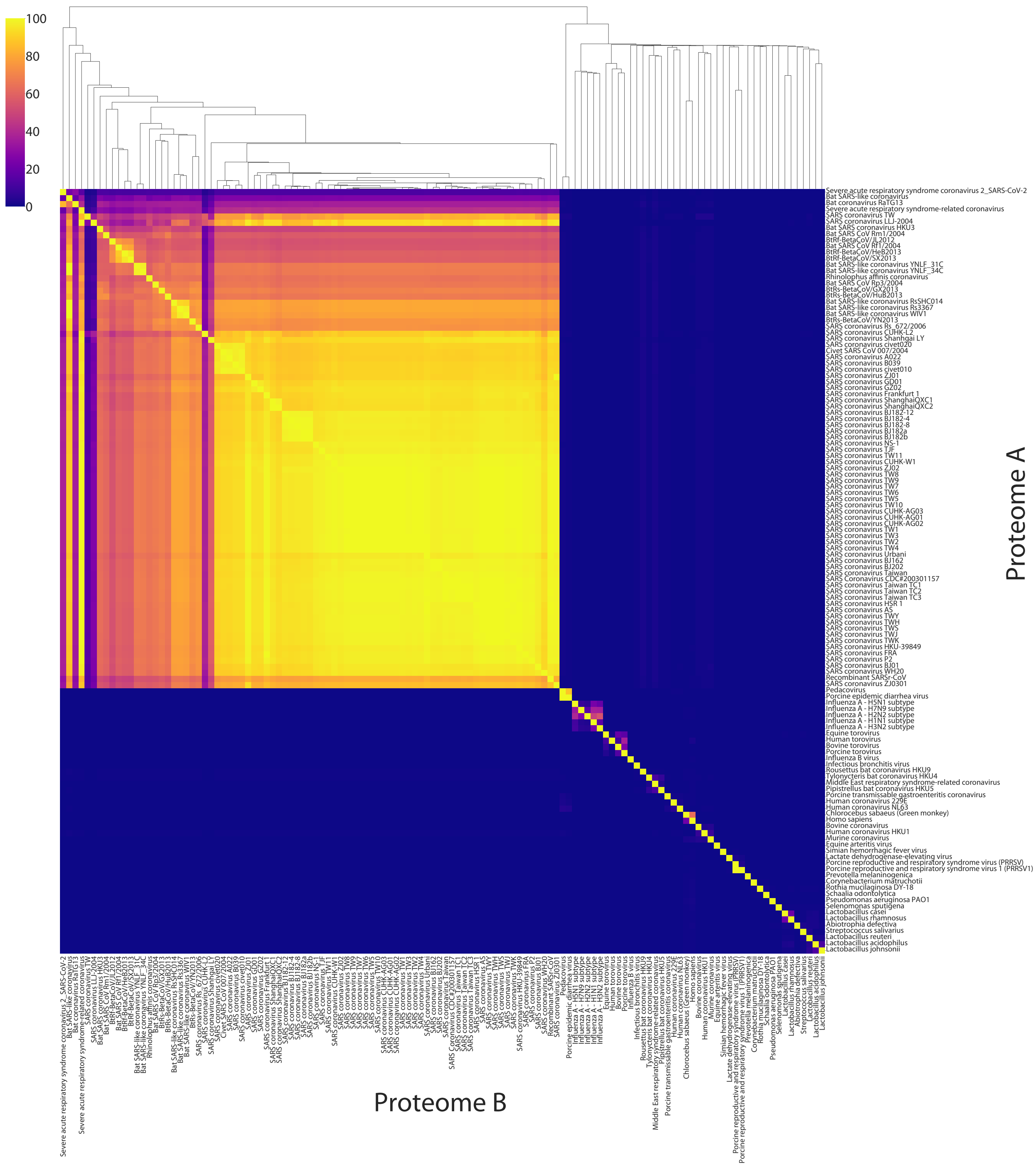


**SI Figure 4.** Heatmap with cluster dendrogram displaying the individual percents of shared tryptic peptides across all taxa currently in the METATRYP Coronavirus database (see SI Figure 5 for annotations).

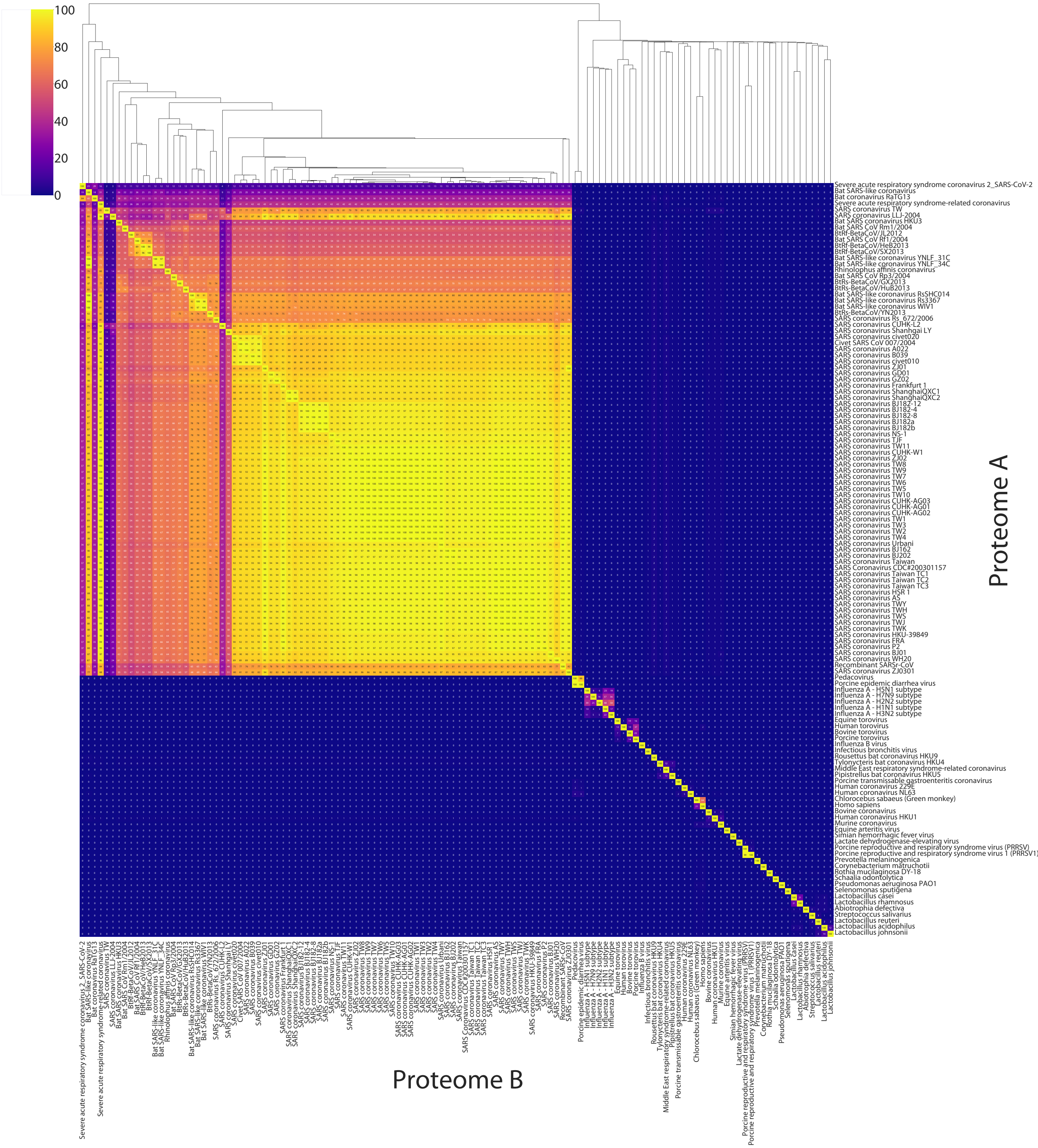


**SI Figure 5.** Heatmap with cluster dendrogram displaying the individual percents of shared tryptic peptides across all taxa currently in the METATRYP Coronavirus database.

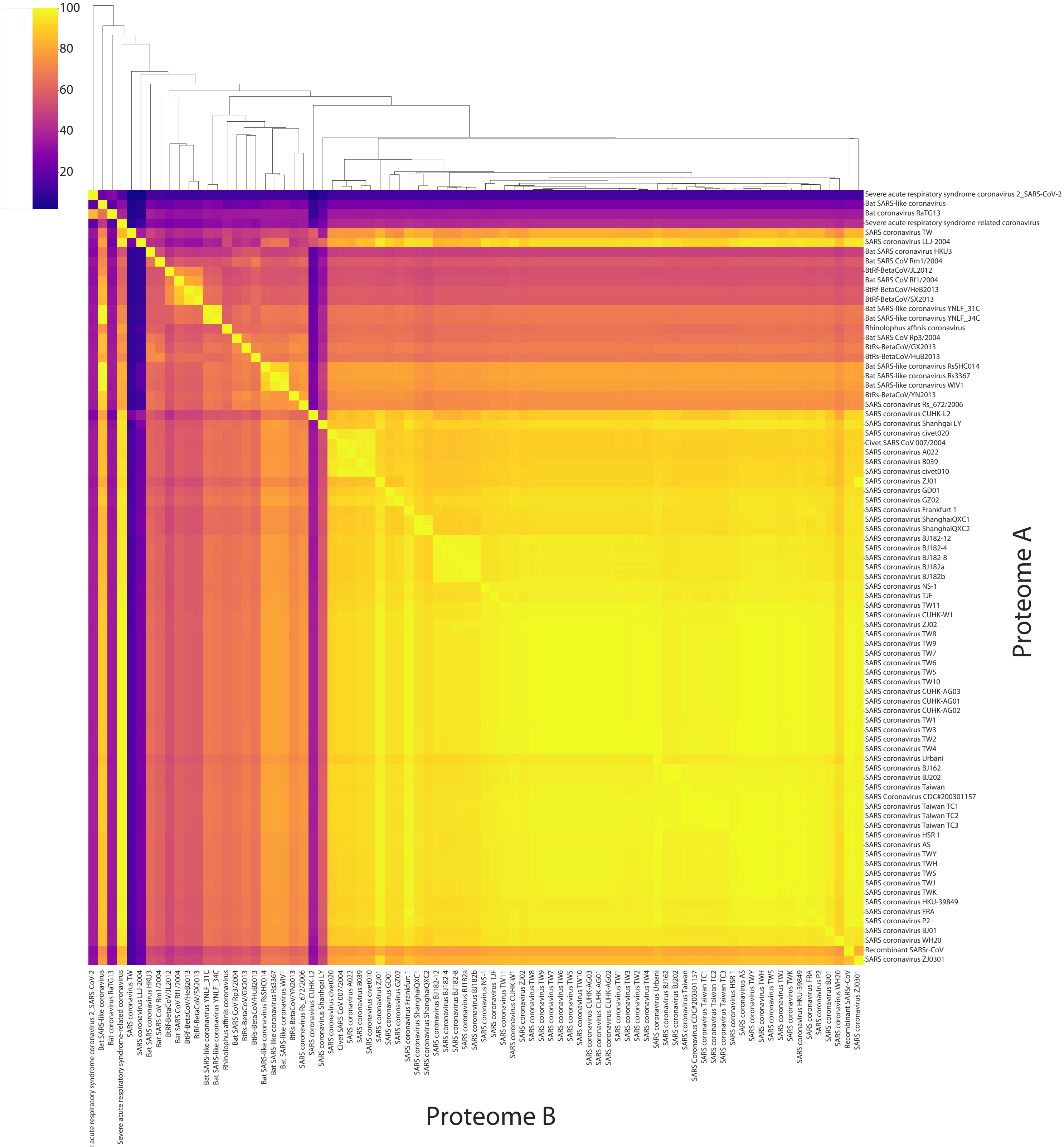


**SI Figure 6.** Heatmap with cluster dendrogram displaying the individual percents of shared tryptic peptides across all the Severe acute respiratory syndrome (SARS) related taxa currently in the METATRYP Coronavirus database (see SI Figure 7 for annotations).

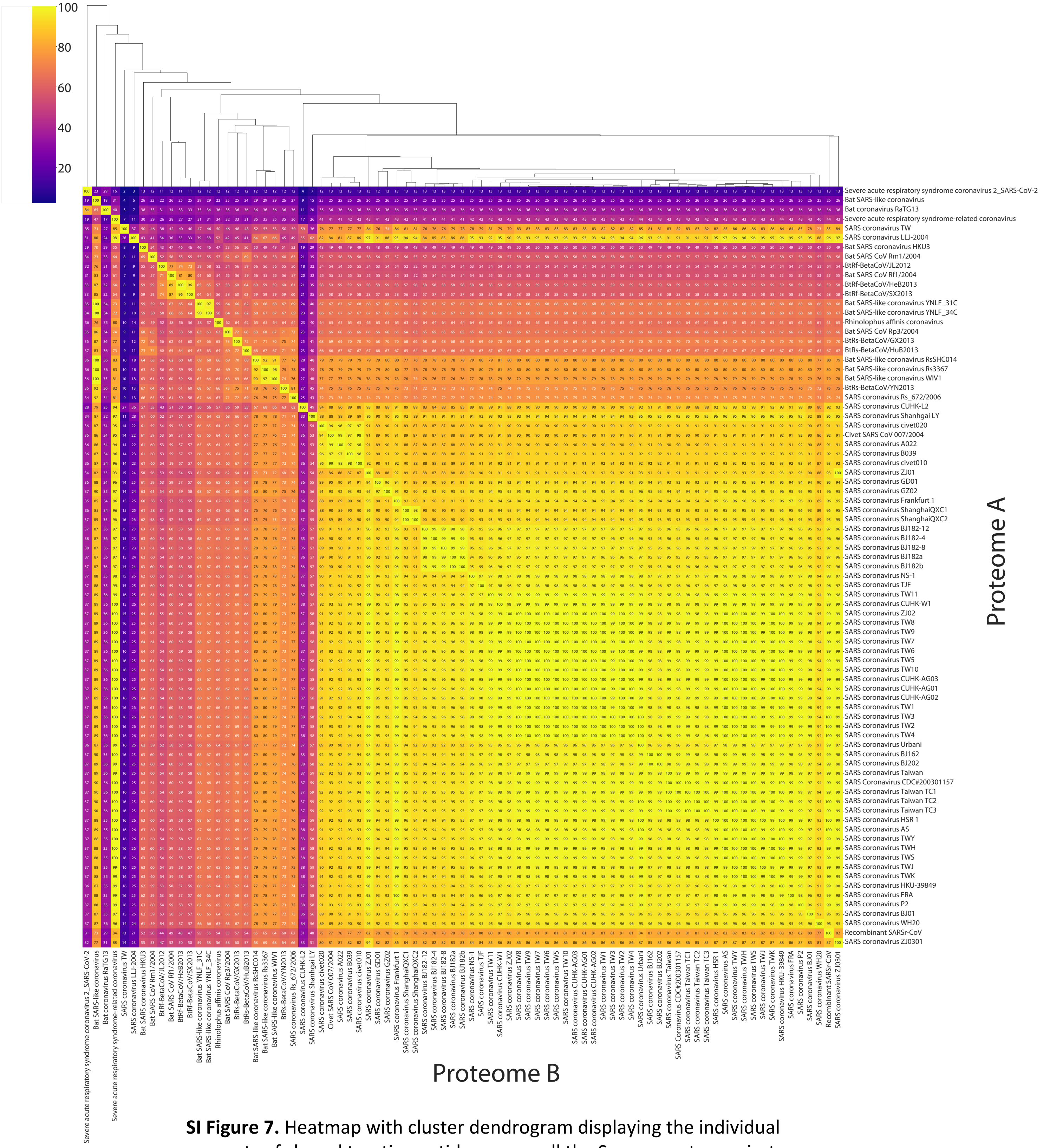


**SI Figure 7.** Heatmap with cluster dendrogram displaying the individual percents of shared tryptic peptides across all the Severe acute respiratory syndrome (SARS) related taxa currently in the METATRYP Coronavirus database.
